# Nuclear Factor Y controls epithelial growth by regulating mTOR in the Drosophila midgut

**DOI:** 10.64898/2026.01.30.702747

**Authors:** Tetiana Strutynska, Onur Deniz, Jaakko Mattila

**Affiliations:** Faculty of Biological and Environmental Sciences, University of Helsinki, Helsinki 00790, Finland; Institute of Biotechnology, University of Helsinki, Helsinki 00790, Finland

**Keywords:** Drosophila midgut, tissue growth, NF-Y, mTOR, nutrient sensing, intestinal stem cell

## Abstract

The intestinal epithelial lining is highly dynamic, with size and cellular composition adapting to nutrient status. This requires regulation of intestinal stem cell (ISC) proliferation and enterocyte (EC) size. How the absorptive area of the intestine matches physiological nutrient conditions remains unclear. Here, we show that the Nuclear Factor Y (NF-Y) transcription factor plays a role in this process. NF-Y loss-of-function (LOF) in ISCs led to high proliferation and cell growth, a phenotype influenced by dietary nutrients. NF-Y LOF also increased nutrient metabolism, shown by more mitochondria and larger lipid droplets in progenitors. Mechanistically, NF-Y restrains mTOR complex 1 (mTORC1) activity in ISC by controlling transcription of mTORC1 signaling components such as Iml1 and Sestrin. Overall, our results demonstrate that NF-Y limits excessive intestinal epithelial growth under nutrient-rich conditions.

**Highlights:**

- NF-Y is a cell-autonomous regulator of ISC activity
- NF-Y restricts ISC proliferation and epithelial growth under nutrient-rich condition
- NF-Y regulates mitochondrial biogenesis and lipid storage in progenitors
- NF-Y limits mTORC1 activity in progenitors

## INTRODUCTION

The size of the intestinal epithelial lining is dynamically regulated to match the animal’s digestive and metabolic needs. For example, the *Drosophila* midgut, the counterpart of the mammalian small intestine, is highly adaptive to nutritional changes. When calorie-restricted, intestinal stem cell (ISC) proliferation is reduced, and the midgut absorptive area decreases due to enterocyte (EC) loss and reduced cell size. Upon feeding, the proliferation and differentiation of ISCs increase, and the ECs enlarge, leading to an increase in absorptive epithelial area (1–3). Although the cellular processes of midgut adaptation have been described in fine detail, the mechanisms underlying the regulation of optimal epithelial size under physiological conditions remain poorly understood.

Adaptive regulation of intestinal absorptive area is mediated by nutrient-sensing pathways, including the amino acid-sensing mTOR complex 1 (mTORC1) signaling pathway (1, 3). In ISCs, mTORC1 regulates anabolic metabolism, enabling cell growth and division, while in progenitors, mTORC1 activity is linked to lineage choice and differentiation (2, 4–6). In niche cells, mTORC1 functions cell non-autonomously to regulate the proliferative activity of stem cells (7). Thus, mTORC1 has a profound and complex role in adaptive regulation of the intestinal epithelium. Unrestricted mTORC1 activation can have catastrophic consequences for cells (8). Therefore, feedback mechanisms that limit mTORC1 activity have evolved. These mechanisms ensure optimal mTORC1 activity, balancing cellular growth and division to available nutrients, growth factors, and energy (9). However, how mTORC1 activity is balanced to achieve optimal epithelial size in physiological conditions, especially in complex tissues such as the *Drosophila* midgut, remain enigmatic.

Nuclear Factor Y (NF-Y) is a conserved and pleiotropic trimeric transcription factor composed of the subunits NF-YA, NF-YB, and NF-YC (10). NF-Y regulates gene expression by binding to the CCAAT box via NF-YA, the DNA-binding subunit of the complex (11). NF-Y plays an essential role in early zygotic development in mammals and is a key regulator of cell-cycle progression in proliferating cells, including hematopoietic and muscle stem cells (12–16). Despite these established functions in development and the cell cycle, little is known about the physiological roles of NF-Y transcription factors. Here, we report an unexpected role for NF-Y in regulating *Drosophila* midgut epithelial growth. Specifically, we show that NF-Y inhibits ISC proliferation, mitochondrial biogenesis, and lipid accumulation in progenitors under nutrient-rich conditions, thereby protecting the intestinal epithelial lining from excessive growth and epithelial disruption. Mechanistically, NF-Y limits mTORC1-mediated ISC activation by regulating transcription of mTORC1 signaling components, such as Iml1 and Sestrin. Through this mechanism, we identify NF-Y as a key regulator of intestinal epithelial cells under conditions of high growth demand.

## RESULTS

### Nuclear Factor Y restricts intestinal stem cell activity cell autonomously

By systematically analyzing transcriptional regulators, we discovered a novel role for Nuclear Factor Y subunit Alpha (NF-YA) in the adult *Drosophila* midgut. When knocked down in female flies using the intestinal stem cell (ISC)-specific driver esg-Gal4^ts^, Su(H)-Gal80 (hereafter referred to as ISC-Gal4) driven RNA interference (RNAi), we observed a substantial expansion of the YFP-expressing cell pool compared with control animals (Figure 1A). The YFP-positive cells accumulate in a highly region-specific manner in regions R2, R4, and R5, whereas regions R1, R3, and the flanking border regions are unaffected (Figure 1A). This data was further confirmed using an independent RNAi line, which produced a similar region-specific expansion of the YFP-positive cells (Figure S1A). Quantification of the phenotype showed significantly increased total and YFP-positive cell numbers (Figure 1B, C). Staining with NF-YA antibodies confirmed the loss of NF-YA protein from cells expressing the RNAi reagent (Figure 1A). The increase in cell numbers suggests an elevated rate of ISC proliferation in response to NF-YA loss-of-function (LOF). To test this possibility, we used 5-Ethynyl-2’-deoxyuridine (EdU) to label cells with newly synthesized DNA. In control animals, a 16h EdU pulse labels YFP-negative enteroblasts (EBs) and small YFP-positive ISCs that have begun endoreduplication or S-phase, respectively (Figure S1B). NF-YA knockdown by the ISC-Gal4 driver resulted in a significant increase in the number of EdU-positive ISCs compared with controls, indicating increased proliferative activity (Figure 1D & E).

**Figure 1.**
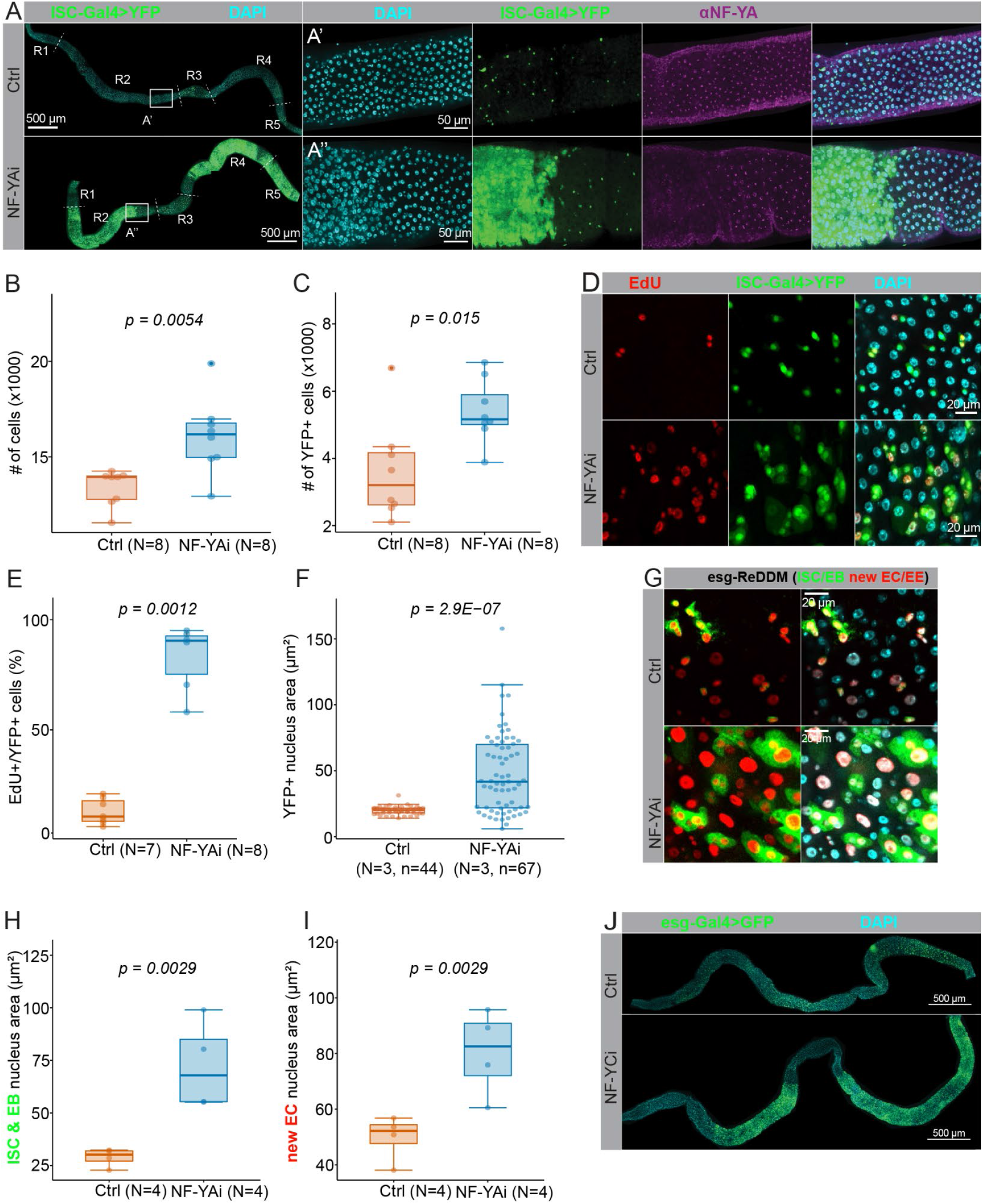
NF-Y restricts intestinal stem cell proliferation and progenitor growth. (**A**) Representative images of esg-Gal4^ts^, Su(H)GBE-Gal80 > NF-YA RNAi (KK) and control female midguts. In **A’ & A’’** are shown the R3-R4 border regions stained by the α-NF-YA antibodies (magenta). Flies were kept at +18°C for 3 days in minimal 2% sucrose media, shifted to a permissive temperature (+29°C) for 3 days to induce RNAi, and then kept on a holidic diet for an additional 7 days. (**B & C**) Quantification of midgut total cell numbers (B) and YFP-positive cells (C) from the experiment depicted in A. (**D**) Representative images of esg-Gal4^ts^, Su(H)GBE-Gal80 > NF-YA RNAi and control female midguts stained by EdU. Images from the R4b region after 3 days on a holidic diet followed by a 16h pulse on a holidic diet supplemented with EdU. (**E & F**) Quantification of the number of EdU-positive ISCs (E) and the nuclear area of YFP-positive cells (F) from the experiment depicted in D. (**G**) Representative images of the esg-Reddm NF-YA RNAi and control female flies from the R4b region after tracing for 7 days on a holidic diet. ISCs/EBs in green & red and newly formed ECs in red. (**H & I**) Quantification of nuclear area of ISCs/EBs (H), and newly formed ECs (I) from the experiment depicted in G. (**J**) Representative images of esg-GAL4^ts^ > NF-YC RNAi and control female midguts. P-values shown in (B), (C), (E), (F), (H), and (I) were obtained by the Wilcoxon rank-sum test.

NF-YA localizes to the nucleus of ISCs as well as in the polyploid enterocytes (ECs) (Figure 1A & S1C). Thus, we asked whether NF-YA regulates ISCs cell autonomously or through non-cell-autonomous function from the EB/EC population. To this end, we used the Su(H)-Gal4 and Myo1A-Gal4 drivers to knock down NF-YA in the EBs and ECs, respectively. Knocking down NF-YA in the EB/EC population did not lead to noticeable expansion of the progenitor cell pool (Figure S1D & S1E). Interestingly, NF-YA knockdown in ISCs led to a broad size distribution of YFP-positive cells, compared with the uniformly small cell sizes in control animals (Figure 1F). As a surrogate for cell size, we measured the maximum nuclear cross-sectional area, which we previously showed to correlate with cell size in the adult midgut (2). In control animals, ISC-Gal4-driven YFP expression is restricted to diploid ISCs by the EB-specific expression of Gal80 (Su(H)-Gal80). However, following NF-YA loss, many YFP-positive cells enlarge, suggesting rapid growth and retention of the YFP signal. To determine whether these cells are progenitors or mature ECs, we stained midguts with antibodies against HRP, a known progenitor marker of the *Drosophila* midgut (17).

Interestingly, most enlarged YFP-positive cells are also anti-HRP-positive, indicating that, despite their large size, these cells are undifferentiated progenitors (Figure S1F). To ask whether the enlarged progenitors also produce larger progeny, we employed the esg-ReDDM (Repressible Dual Differential stability cell Marker) lineage-tracing method to distinguish between esg-expressing progenitors and fully matured progeny (18). The esg-ReDDM utilizes the differing stabilities of the short-lived membrane-targeted GFP, which acts as a precise temporal marker of esg-Gal4 activity, and the long-lived histone 2B-tagged RFP, enabling tracking of newly differentiated progeny. Our analysis shows that NF-YA knockdown by the esg-gal4 driver results in a significant increase in progenitor and progeny size compared with controls (Figure 1H-I). Thus, based on our findings, we conclude that NF-YA functions in ISCs to inhibit progenitor growth and progeny size.

NF-YA is the sequence-binding motif of the conserved heterotrimeric Nuclear Factor Y (NF-Y) transcription factor, which forms a functional trimer with the NF-YB and NF-YC subunits (10). To confirm that the NF-YA LOF phenotype in ISCs results from the disruption of the NF-Y complex, we knocked down NF-YC using esg-Gal4-driven RNAi. Indeed, knockdowns of NF-YA and NF-YC produced identical, region-specific phenotypes, confirming that disruption of the functional NF-Y trimer underlies these phenotypes (Figure 1J). In conclusion, our results show that NF-Y has a cell-autonomous role in restricting ISC proliferation and progenitor growth in a region-specific manner.

### Disrupting NF-Y in ISCs leads to epithelial growth and reduced lifespan

The increase in midgut cell number and cell size upon NF-Y disruption in ISCs suggests an expansion of tissue size. However, measuring the lengths of midgut regions in animals with ISC-Gal4-driven NF-YA knockdown under nutrient-rich conditions (holidic diet (19)) revealed no significant differences in regional midgut lengths compared with controls (Figure 2A). We then asked how the midgut epithelial structure is altered by the increase in cell number and cell size in these animals. Interestingly, the apico-basal epithelial structure in the midguts of NF-YA knockdown animals showed dramatic cellular multilayering, compared with the single-layer organization in control midguts (Figure 2B). In addition, knockdown of NF-YA resulted in cellular delamination from the epithelium into the lumen (Figure 2B). Thus, under high-nutrient conditions, the excess, oversized cells in the ISC-Gal4-driven NF-YA knockdown are multilayering and delaminating from the epithelium. The phenotype became more severe with age, resulting in a strikingly narrowed gut lumen (Figure 2B). To test the physiological relevance of the distorted epithelial structure, we monitored the lifespan of flies with NF-YA knockdown driven by the ISC-Gal4 driver. Indeed, these animals showed a significantly shorter lifespan than controls (Figure 2C). Thus, NF-Y is important for maintaining optimal epithelial size, and its disruption can reduce animal survival. Interestingly, while searching for physiological conditions that adjust the NF-YA LOF phenotype, we observed that when flies were maintained on a minimal 2% sucrose diet, the epithelial disruption was completely rescued (Figure 1K-N). Taken together, our results show that NF-Y is an obligate inhibitor of ISC activation and epithelial growth under nutrient-rich conditions.

**Figure 2.**
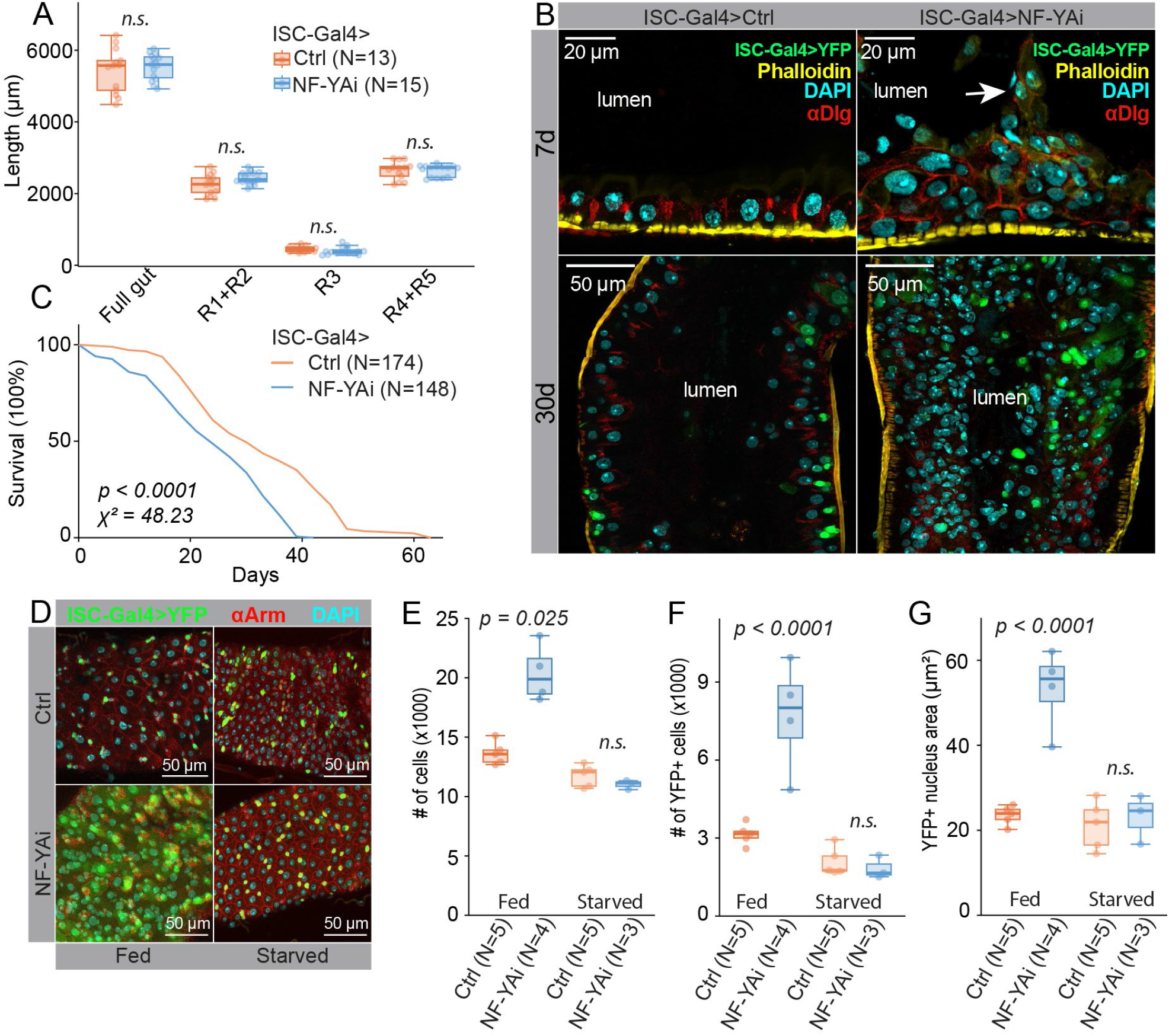
NF-Y regulates epithelial growth in nutrient-rich conditions. (**A**) Quantification of esg-Gal4^ts^, Su(H)GBE-Gal80 > NF-YA RNAi (KK) and control midgut region lengths. (**B**) Representative images of esg-Gal4^ts^, Su(H)GBE-Gal80 > NF-YA RNAi (KK) and control midgut cross-sections from the R4b region immunostained with α-Dlg (septate junctions) and phalloidin (F-actin). Flies kept for 7 days (young) or 30 days (old) on a holidic diet at +29°C. An arrow points to a group of cells delaminating from the epithelium. (**C**) Quantification of esg-Gal4^ts^, Su(H)GBE-Gal80 > NF-YA RNAi (KK) and control virgin female survival at +29°C. (**D**) Representative images of esg-Gal4^ts^, Su(H)GBE-Gal80 > NF-YA RNAi (KK) and control midguts from the R4b region from flies kept either on a holidic diet (fed) or 2% sucrose (starved) and immunostained with α-Armadillo antibodies. (**E-G**) Quantification of total cell numbers (E), YFP+ cell numbers (F), and YFP+ nuclear area (G) from the experiment depicted in (D). Significances in (A) were obtained by two-way ANOVA followed by Tukey’s test. P-value and chi-square value in (C) were obtained by the Log-rank (Mantel-Cox) test. P-value in (E) was obtained by Welch ANOVA followed by the Games-Howell test. P-values in (F) and (G) were obtained by two-way ANOVA followed by Tukey’s test.

### NF-Y regulates mitochondrial biogenesis and lipid storage in ISCs

To obtain a global view of the biological processes regulated by NF-Y, we performed mRNA sequencing in NF-YA-deficient ISCs. To restrict our analysis to early regulatory events, we isolated mRNA from ISCs after 3 days of RNA interference induction. At this time, the NF-YA knockdown phenotype is not yet fully penetrant, suggesting that the observed changes in gene expression are directly regulated by NF-Y (Figure S2A). To isolate ISCs, we used fluorescence-activated cell sorting of YFP-marked ISCs. From the dataset, we performed differential gene expression analysis (DEG) and identified 850 upregulated and 989 downregulated genes, including NF-YA among the ten most downregulated genes (Figure 3A, Table S1). Gene set enrichment analysis (GSEA) revealed that the most affected biological processes are mitochondrial translation & transcription and the cell cycle (Figure S2B). Indeed, the expression of several key cell cycle drivers, such as *cyclin B, cyclin E, cdk1*, and *PCNA*, was significantly upregulated upon NF-YA LOF in ISCs (Figure S2C-F). The elevated transcription of those genes is consistent with the hyperproliferation of ISCs resulting from NF-Y disruption. In addition, the expression of several components of the inner (Tim proteins) and outer (Tom proteins) membrane translocase complexes of the mitochondrion was significantly upregulated in NF-YA LOF ISCs, including *Tom20, Tom70, Tim14*, and *Tim17b* (Figure 3B-E). These data suggest an increased number or volume of mitochondria in NF-YA LOF ISCs. In line with this, several genes involved in mitochondrial metabolism, including those in the TCA cycle and oxidative phosphorylation, were also significantly upregulated by NF-Y disruption (Figure S2G). Finally, the expression of genes promoting lipid droplet formation, such as *seipin* and *schlank*, as well as the transcription factor *sugarbabe*, which regulates genes involved in *de novo* lipogenesis, was upregulated upon NF-YA LOF in ISCs (Figure S2H-J). In conclusion, NF-Y regulates the expression of genes involved in the cell cycle, mitochondrial metabolism, and lipid synthesis & storage in ISCs.

**Figure 3.**
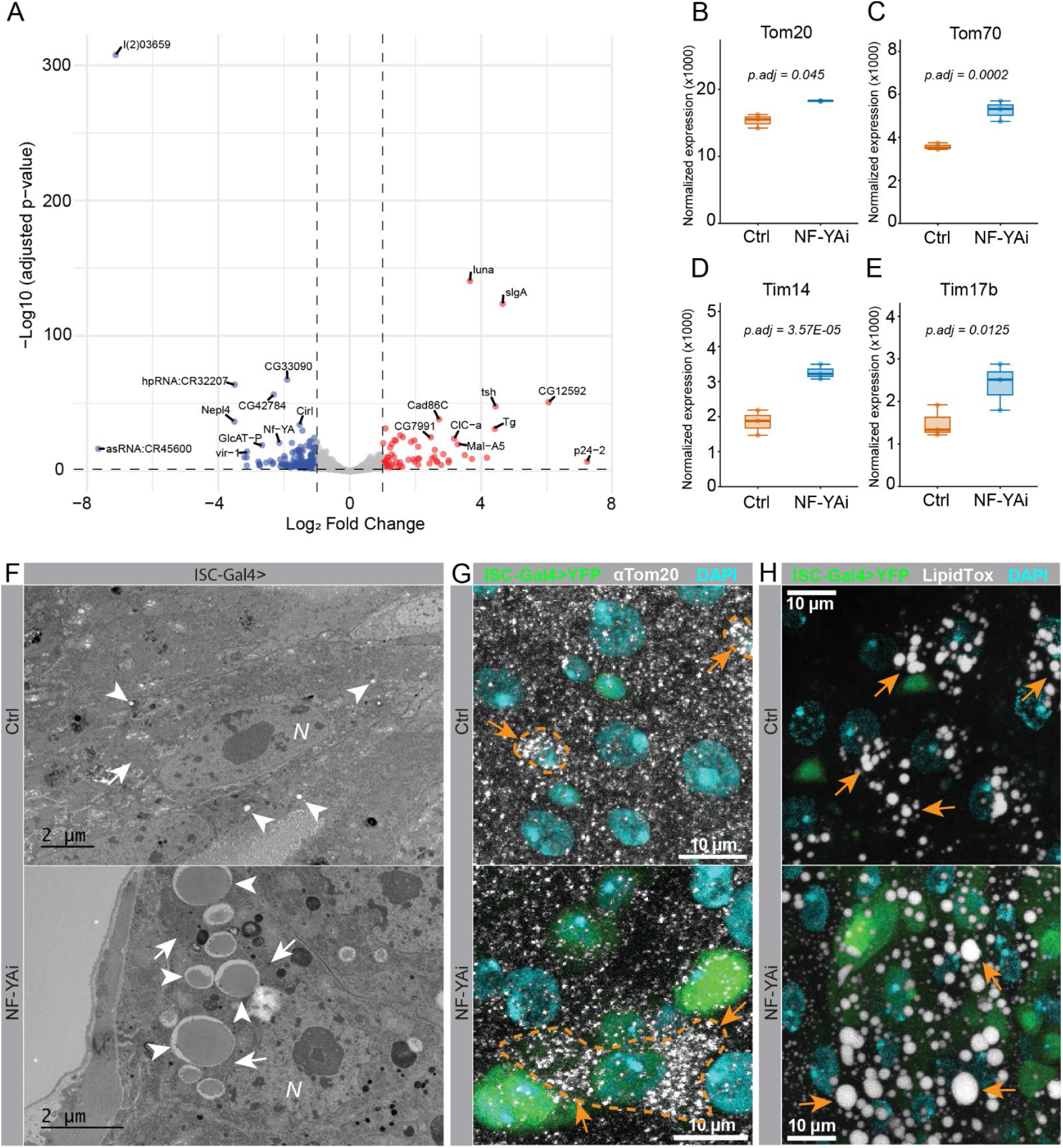
NF-Y regulates mitochondrial biogenesis and lipid storage. (**A**) Volcano plot representing differentially expressed genes between esg-Gal4^ts^, Su(H)GBE-Gal80 > NF-YA RNAi (KK), and control ISCs. Significant changes (p<0.05 and log2 fold changes>±1) are shown in red (upregulated in NF-YA RNAi) and blue (downregulated in NF-YA RNAi). The 10 most up- and downregulated genes are annotated on the plot. (**B-E**) Normalized mRNA expressions of *Tom20* (B), *Tom70* (C), *Tim14* (D), and *Tim17b* (E). (**F**) Electron micrograph from 30d-old esg-Gal4^ts^, Su(H)GBE-Gal80 > NF-YA RNAi (KK) and control midguts from the R5 region. Arrows point to lipid droplets, and arrowheads point to mitochondria. (**G**) Representative images of 7d-old esg-Gal4^ts^, Su(H)GBE-Gal80 > NF-YA RNAi (KK) and control midguts from the R4b region immunostained with α-Tom20 antibodies. Arrows point to α-Tom20 puncta. (**H**) Representative images of 7d-old esg-Gal4^ts^, Su(H)GBE-Gal80 > NF-YA RNAi (KK) and control midguts from the R4b stained with LipidTox. Arrows point to lipid droplets. Adjusted p-values in A-E were obtained by the DESeq2 package with Benjamini-Hochberg correction.

To directly assess NF-Y’s role in regulating cellular processes, we examined subcellular structures by transmission electron microscopy (TEM). We analyzed midguts from ISC-Gal4-driven NF-YA RNAi animals at 30 days of age, ensuring that the epithelium had fully turned over by progenitors derived from NF-Y LOF ISCs. In line with the observed gene expression changes, we observed increased mitochondrial number and size upon NF-Y disruption compared with controls (Figure 3F). Strikingly, the mitochondria surrounded greatly enlarged lipid droplets (Figure 3F). To exclude the possibility that the observed cellular changes were specific to old age, we immunostained midguts from seven-day-old animals with antibodies against the mitochondrial import receptor subunit Tom20 and the neutral lipid stain LipidTox. Consistent with the TEM analysis, we observed an increased number of anti-Tom20 immunostained puncta and larger lipid droplets in ISC-Gal4-driven NF-YA RNAi cells compared with wild-type cells (Figure 3G & H). Taken together, our results show that NF-Y is an essential regulator of mitochondrial biogenesis and lipid storage in ISCs and their progeny.

### NF-Y acts as a metabolic and proliferative brake through inhibiting mTORC1 signaling

To understand how NF-Y regulates ISCs under nutrient-rich conditions, we identified NF-YA-regulated genes involved in nutrient sensing in our dataset. Indeed, we found several prominent negative regulators of the mTOR complex 1 (mTORC1) among the genes downregulated upon ISC-Gal4-driven NF-YA RNAi, including *pras40, sestrin, iml1*, and *scylla*/*charybdis* (Figure 4A). In addition, the expression of transcriptional effectors *reptor* and *reptor-bp* was also downregulated in these cells (Figure 4A). Attenuation of Reptor is needed for the activation of the mTORC1-mediated transcriptional response (20). These results suggest that NF-Y inhibits mTORC1 signaling in ISCs. To directly test this hypothesis, we stained midguts with antibodies against the phosphorylated form of the mTORC1 target initiation factor 4E–binding protein (p4EBP). Indeed, we observed very high p4EBP levels in the NF-YA LOF progenitors, indicating elevated mTORC1 activity relative to controls (Figure 4B). To test whether the elevated mTORC1 activity is sufficient to drive the NF-YA LOF-induced ISC proliferation, cell size increase, and epithelial disruption, we inhibited mTORC1 through rapamycin feeding or genetically by knocking down the Regulatory-Associated protein of mTOR (Raptor) (21). Strikingly, we found that both rapamycin feeding and ISC-Gal4-driven knockdown of Raptor rescued the NF-YA LOF phenotype in ISCs (Figure 4C-F, Figure S3). These results show that NF-Y is an essential inhibitor of mTORC1 signaling in ISCs, thereby preventing overproliferation, progenitor growth, and epithelial disruption in nutrient-rich conditions.

**Figure 4.**
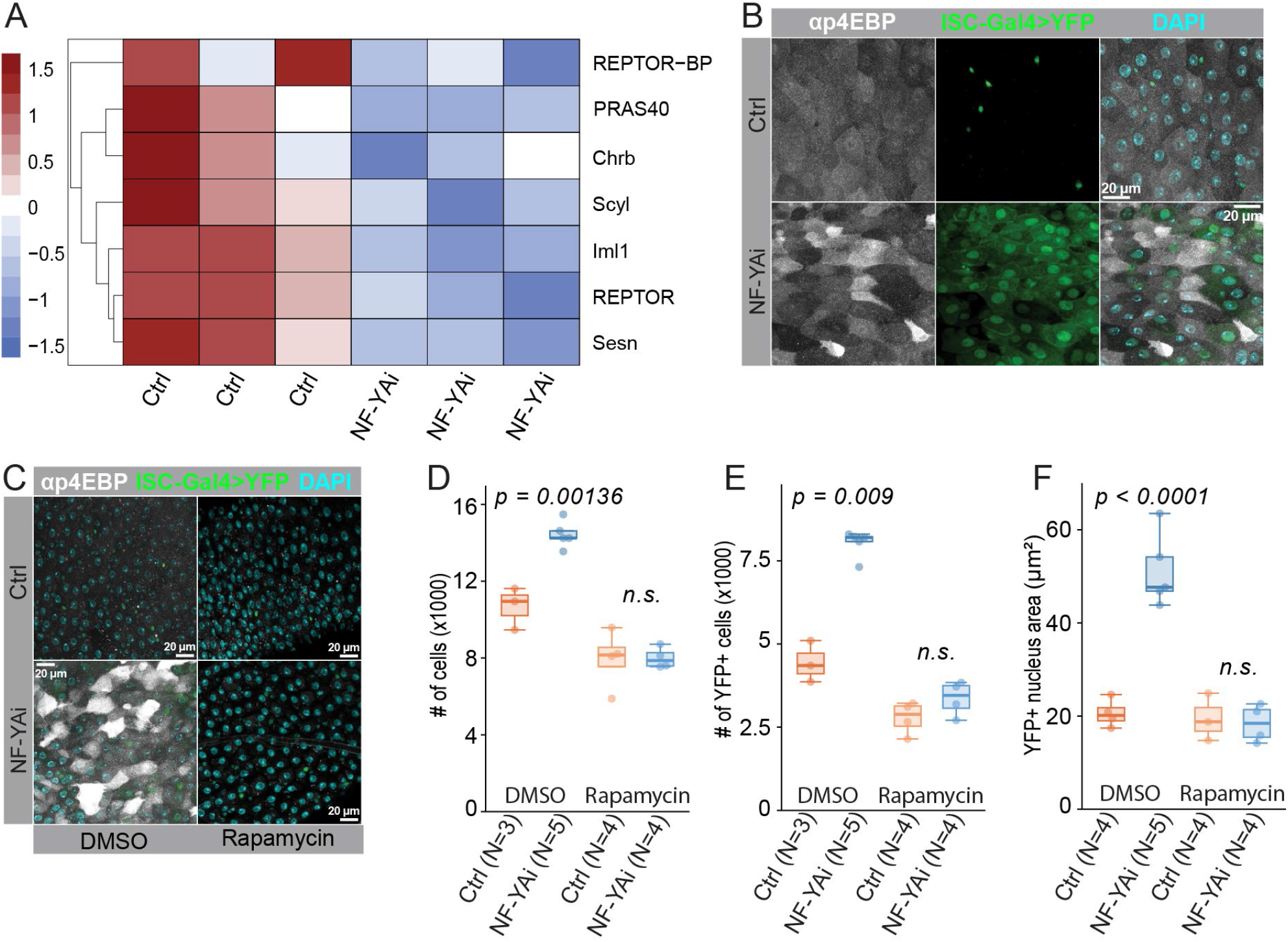
NF-Y regulates proliferation and cell growth via mTORC1. (**A**) Heatmap of differentially expressed genes involved in the mTORC1 signaling pathway (GO database) from the mRNA sequencing experiment depicted in Figure 3A. (**B**) Representative images of esg-Gal4^ts^, Su(H)GBE-Gal80 > NF-YA RNAi (KK) and control female midguts from the R4b region immunostained with α-p4EBP antibodies. Flies were kept at +29°C for 4 days. (**C**) Representative images of esg-Gal4^ts^, Su(H)GBE-Gal80 > NF-YA RNAi (KK) and control female midguts from the R4b region kept on a holidic diet supplemented with either DMSO (control conditions) or 200 μM rapamycin. (**D-F**) Quantification of total cell numbers (D), YFP+ cell numbers (E), and YFP+ nuclear area (F) from the experiment depicted in (C). P-values for (D) and (E) were obtained by two-way ANOVA followed by Tukey’s test. P-value in (F) was obtained by Welch analysis of variance (ANOVA) followed by the Games-Howell test.

## DISCUSSION

Our study elucidates a new and unexpected role for NF-Y in *Drosophila* ISCs. Under nutrient-rich conditions, NF-Y limits ISC proliferation and enterocyte size, reducing intestinal epithelial growth. Mechanistically, NF-Y regulates genes involved in the cell cycle, mitochondrial metabolism, lipid biosynthesis, and mTORC1 signaling. Thus, NF-Y acts as a metabolic brake, protecting the intestinal epithelium from excess growth in nutrient-rich conditions.

NF-Y is a highly conserved trimeric eukaryotic transcription factor. It consists of the DNA-binding NF-YA and the regulatory subunits NF-YB and NF-YC. NF-Y plays key roles in many cell types and cellular processes. These range from early zygotic development to metabolism, the cell cycle, and apoptosis (10). Our results show that NF-Y has dual functions in ISCs. First, it restricts ISC proliferation by regulating the expression of key cell cycle regulators. Second, it reduces progenitor growth and the size of mature enterocytes. Interestingly, NF-YA knockdown in enteroblasts—the progenitors of enterocytes—did not increase cell growth in our conditions. This implies that NF-Y’s growth-restricting function in progenitors occurs already in ISCs. How does NF-Y mediate its function in ISCs and lead to increased progenitor growth and progeny size? A possible mechanism is through chromatin remodeling in ISCs. NF-Y interacts with corepressors and co-activators in a cell- and context-specific manner, acting as a platform for chromatin remodeling proteins, integrating chromatin structure with transcriptional regulation (22). For example, NF-Y was previously shown to regulate the chromatin landscape during early zygotic development. Depletion of maternally derived NF-YA in a mouse 2-cell embryo led to loss of open chromatin at promoter sites and developmental arrest (23). Future work is needed to resolve the exact role of NF-Y in chromatin remodeling in *Drosophila* ISCs. For example, identifying NF-Y’s binding partners in this setting could clarify its regulation on progenitor growth.

Beyond its role in early zygotic development, NF-Y has been linked to the regulation of glucose and fatty acid metabolism in various cell types, including pancreatic β-cells, cardiomyocytes, hepatocytes, and adipocytes (24–27). In *Drosophila* ISCs, NF-Y represses mitochondrial metabolism and lipid accumulation by regulating gene expressions involved in these processes. In addition, NF-Y restricts mTORC1 activity by regulating the expression of several pathway components. Regulation of mTORC1 signaling is a pivotal cellular mechanism for adjusting growth and anabolic metabolism to prevailing nutrient and energy availability (28). The finding that inhibiting mTORC1 activity in ISCs completely rescued the NF-Y LOF phenotype suggests that NF-Y’s metabolic role is largely mediated by mTORC1. Thus, it would be interesting to know whether the same is true in other cellular contexts.

In ISCs, mTORC1 signaling primes the stem cells to differentiate toward the absorptive lineage, which, during differentiation, undergoes a massive increase in cellular size (2, 29). Constitutive activation of the mTORC1 pathway in ISCs, through loss-of-function of the tuberous sclerosis complex 1/2, leads to overgrowth, cell cycle arrest, and a gradual loss of ISCs, possibly through delamination (5, 29). Simultaneous activation of a pro-proliferative signal, such as a Notch loss-of-function condition, relieves the cell cycle arrest imposed by hyperactive mTORC1 in ISCs and leads to massive epithelial overgrowth (5, 29). Thus, it is probable that NF-Y, in addition to regulating mTORC1 activity, also controls the activity of a pro-proliferative signal in ISCs. Future studies are needed to identify the full scope of NF-Y-regulated pathway activities in ISCs.

## MATERIALS AND METHODS

### Drosophila stocks and husbandry

Fly stocks used in the study: esg-Gal4, Tub-Gal80^ts^, Su(H)GBE-Gal80, 2xYFP (ISC-Gal4) (30), NF-YA RNAi (VDRC 106132, Bloomington 25991), NF-YC RNAi (Bloomington 58234), Raptor-RNAi (Bloomington 31529), Su(H)-Gal4, UAS-GFP; Tub-Gal80^ts^ (31), MyO1A-Gal4, Tub-Gal80^ts^; UAS-GFP (32), Esg-ReDDM (18). Fly stocks were maintained at +25°C, on nutritional medium containing agar 0.6% (w/v), malt 6.5% (w/v), semolina 3.2% (w/v), baker’s yeast 1.8% (w/v), nipagin 2.4%, and propionic acid 0.7%. In all experiments, flies were reared at +18°C and then transferred to +29°C to inhibit temperature-sensitive Gal80, thereby activating the UAS-GAL4 system in the midgut.

### Dietary treatments

Flies were kept on a chemically defined (holidic) diet (19), referred to as fed condition. For starvation, flies were kept on medium containing agar 0.6% (w/v), sucrose 2% (w/v), nipagin 2.4%, and propionic acid 0.7%. The pH of the starvation medium was adjusted to match the holidic diet (pH=6.8) using 5 M NaOH, thus excluding pH as a variable. For rapamycin feeding, holidic diet was supplemented with 200 μM rapamycin (Thermo Scientific Chemicals, CAS #53123-88-9).

### Lifespan assay

Virgin ISC-Gal4 > NF-YA RNAi and control female flies were collected and kept at +18°C for 3 days for midgut maturation. The flies were kept at a density of 10 animals per vial, containing holidic diet. The flies were then maintained at 29°C for the remainder of the experiment. The number of dead flies was scored, and the flies were transferred to fresh media every 3 days.

### Immunohistochemistry

For immunofluorescence staining, intestines were dissected in phosphate-buffered saline (PBS) and fixed in 8% PFA for 2 hours. The fixed tissue was then washed with 0.1% Triton X-100 in PBS and blocked with 1% BSA for 1 h. Subsequently, intestines were stained with anti-NF-YA (1:400) (gift from Hideki Yoshida, Kyoto Institute of Technology), anti-β-galactosidase (1:400) (MP Biomedicals 0855976-CF), anti-Prospero (1:1000) (Developmental Studies Hybridoma Bank MR1A), anti-Dlg (1:50) (Developmental Studies Hybridoma Bank 4F3), anti-Armadillo (1:20) (Developmental Studies Hybridoma Bank N27A1), anti-p4EBP (1:400) (Cell signaling 2855), anti-Horseradish Peroxidase HRP 1:200 (Jackson ImmunoResearch Laboratories 2338967), anti-TOM20 1:400 (BD Biosciences, 612278) and Phalloidin 1:300 (Invitrogen A12380). Samples were mounted in Vectashield with DAPI (Vector Laboratories).

### Ethynyldeoxyuridine feeding and Click-it-assay

5-ethynyl-2’-deoxyuridine (EdU) (Sigma-Aldrich 95740-26-4) was added to holidic food at a concentration of 0.2 mg/ml. Flies were fed the medium containing EdU for 16 h, then dissected in PBS and fixed for 30 min in 8% PFA. After fixation, samples were washed with 0.1% Triton X-100 in PBS and incubated for 20 min in the click assay cocktail (Baseclick, ClickTech EdU Proliferation Kit for High-throughput Screening). The cocktail solution was removed, and samples were washed in 0.1% Triton X-100 in PBS for 1 h. Subsequently, samples were mounted in Vectashield with DAPI (Vector Laboratories).

### Lipidtox staining

Midguts were dissected in Shields and Sang M3 medium (Sigma-Aldrich S8398) and fixed in 8% PFA for 30 min. The midguts were washed in PBS and stained with HCS LipidTOX™ Deep Red Neutral Lipid Stain 1:400 (Invitrogen H34477) in PBS for 30 min. The midguts were washed in PBS and mounted in Vectashield with DAPI (Vector Laboratories).

### Microscopy and image processing

Fixed and immunostained midguts were mounted between a microscope slide and a coverslip using 0.12 μm spacers. The samples were imaged using the Aurox Clarity spinning-disk confocal or the Leica SP8 confocal microscope. Images were further processed in ImageJ and segmented using Stardist as previously described (33).

### Transmission electron microscopy

Midguts were dissected in PBS and fixed in 1% glutaraldehyde, 2% PFA, 2 mM CaCl2, and 100 mM cacodylate buffer (pH 7.5) for 1 h. After dehydration, the samples were embedded in Epon resin. Samples were imaged on a Jeol JEM-1400 transmission electron microscope.

### FACS and RNA sequencing

The protocols for FACS and RNA sequencing were adapted from (34). Three independent samples from the control and NF-YA RNAi lines, driven by the ISC-Gal4 driver, were processed in parallel. Female flies were kept on holidic diet for 3 days at +29°C, and then the intestines were dissected and processed into cell lysates. After dissociation, cells were filtered through a 40 μm filter and FACS-sorted based on YFP-signal. RNA was extracted using the Arcturus™ PicoPure™ RNA isolation Kit (Thermo Fisher Scientific 3032689). RNA concentration and quality were assessed using Agilent Bioanalyzer, followed by sequencing on the AVITI High Output (two-end reads, length 75 bp) platform. Raw RNA-seq data were processed using nf-core/rnaseq pipeline (v 3.18.0) (35). Reads were trimmed using Trim Galore (v 0.6.10) and subsequently aligned with STAR (v 2.7.11b) to the *D. melanogaster* reference genome (FlyBase r6.62). Salmon (v 1.10.3) was used to obtain transcript-level counts. All processing steps were orchestrated with Nextflow (v 24.10.3) and executed within Singularity containers (v 3.18.0). Heatmaps were generated with tidyheatmaps (v 0.2.1) (https://jbengler.github.io/tidyheatmaps/authors.html#citation). The raw and processed sequencing data are deposited into the GEO repository (GSE317617). To identify significantly enriched pathways, gene set enrichment analysis (GSEA) included KEGG, WikiPathways, Reactome, and Gene Ontology (GO) databases, and was executed using the clusterProfiler package (version 4.12.6) (36) in R/Bioconductor. Pathways with an adjusted p-value < 0.05 with Benjamini–Hochberg correction were considered significantly enriched.

### Statistical analysis

Statistical analysis was performed using R/Bioconductor and GraphPad Prism. Homogeneity of variances was assessed using Levene’s test, and normality was evaluated using the Shapiro–Wilk test. For comparisons between two groups, either an unpaired Student’s t-test or the Wilcoxon rank-sum test was used, depending on variance homogeneity and data normality. For multiple comparisons, data were analyzed using either two-way ANOVA followed by Tukey’s post hoc test, or Welch’s ANOVA followed by the Games–Howell post hoc test. Survival curves were analyzed using the log-rank (Mantel–Cox) test. The exact test, sample numbers, and P-values for each experiment are provided in the figures and figure legends.

## Supporting information

Supplemental images

## ACKNOWLEDGEMENTS

We thank Hideki Yoshida, Kyoto Institute of Technology, for providing the anti-NF-YA antibody and Heini Lassila for technical assistance. Bloomington Drosophila Stock Center and Vienna Drosophila Resource Center are acknowledged for fly stocks. This study was facilitated by the University of Helsinki Drosophila core facility (Hi-Fly), the Light Microscopy Unit (LMU), the Electron Microscopy Unit (EMBI), the FACS unit, and the Sequencing unit, supported by the Biocenter Finland and the Helsinki Institute of Life Science. This work was supported by Research Council of Finland, Sigrid Jusélius Foundation and Jane & Aatos Erkko Foundation (to J.M.).

## AUTHOR CONTRIBUTIONS

T.S.: investigation, project administration, formal analysis, visualization, conceptualization, writing - original draft, writing - review & editing, methodology; O.D.: investigation, formal analysis; J.M.: conceptualization, supervision, funding acquisition, writing - original draft, writing - review & editing, project administration.

