## Supplemental images for "Nuclear Factor Y controls epithelial growth by regulating mTOR in the Drosophila midgut"

Tetiana Strutynska et al.

FIGURE S1

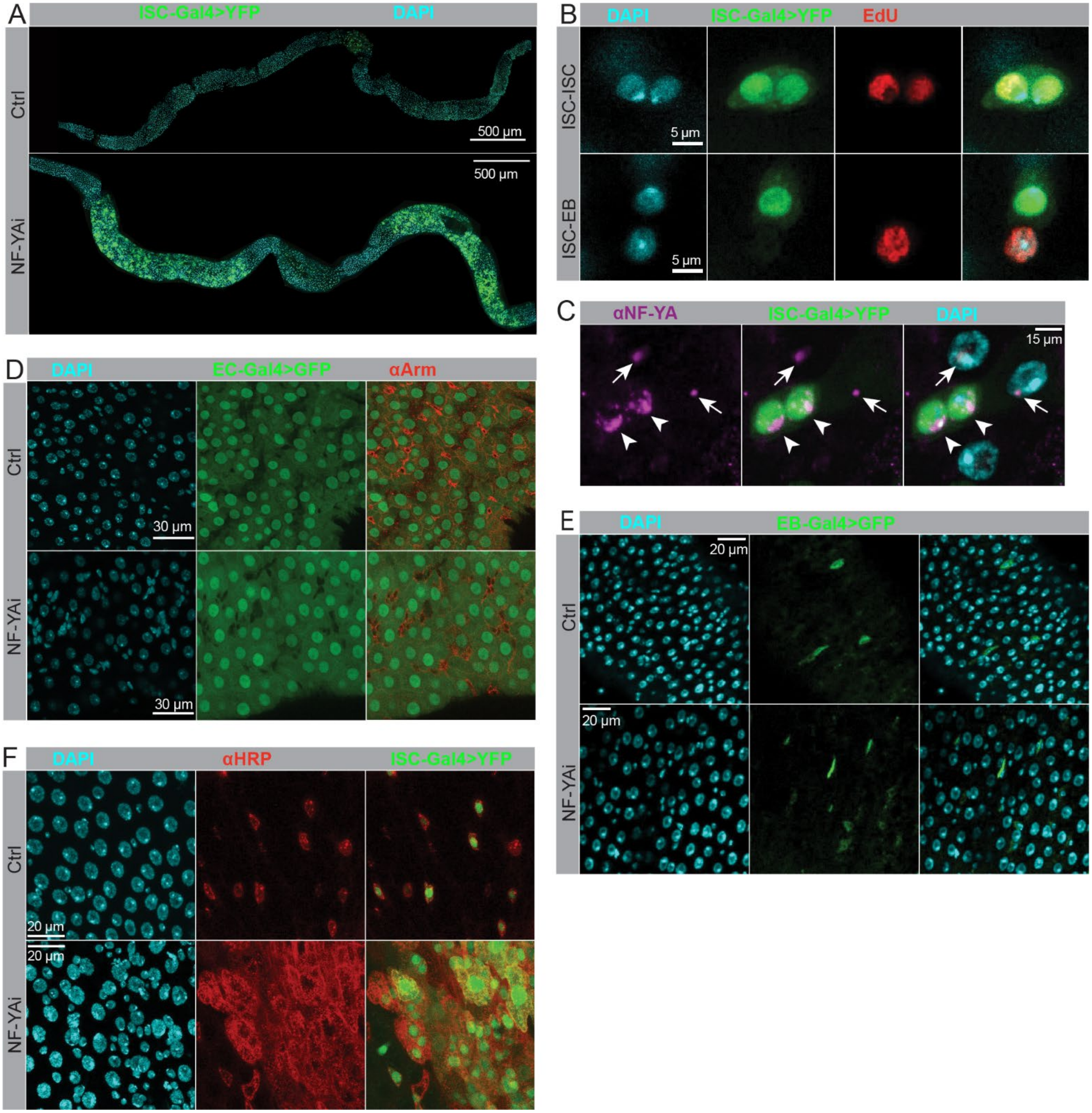


*(Legend on the next page)*

**Figure S1. Related to the main figure 1.**

(**A**) Representative images of esg-Gal4$^{ts}$, Su(H)GBE-Gal80 > NF-YA RNAi (Trip) and control female midguts. (**B**) Representative images of esg-Gal4$^{ts}$, Su(H)GBE-Gal80 > YFP ISC-ISC and ISC-EB cell pairs showing EdU incorporated chromatin. (**C**) Representative images of esg-Gal4$^{ts}$, Su(H)GBE-Gal80 midguts immunostained with α-NF-YA antibodies. Arrowheads point to YFP+ ISCs, and arrows point to ECs. (**D**) Representative images of Myo1A-Gal4$^{ts}$ > NF-YA RNAi (KK) and control midguts from the R4b region immunostained with α-Armadillo antibodies. (**E**) Representative images of Su(H)-Gal4$^{ts}$ > NF-YA RNAi (KK) and control midguts from the R4b region. (**F**) Representative images of esg-Gal4$^{ts}$, Su(H)GBE-Gal80 > NF-YA RNAi (KK) and control female midguts from the R4b region immunostained with α-HRP antibodies.

### FIGURE S2

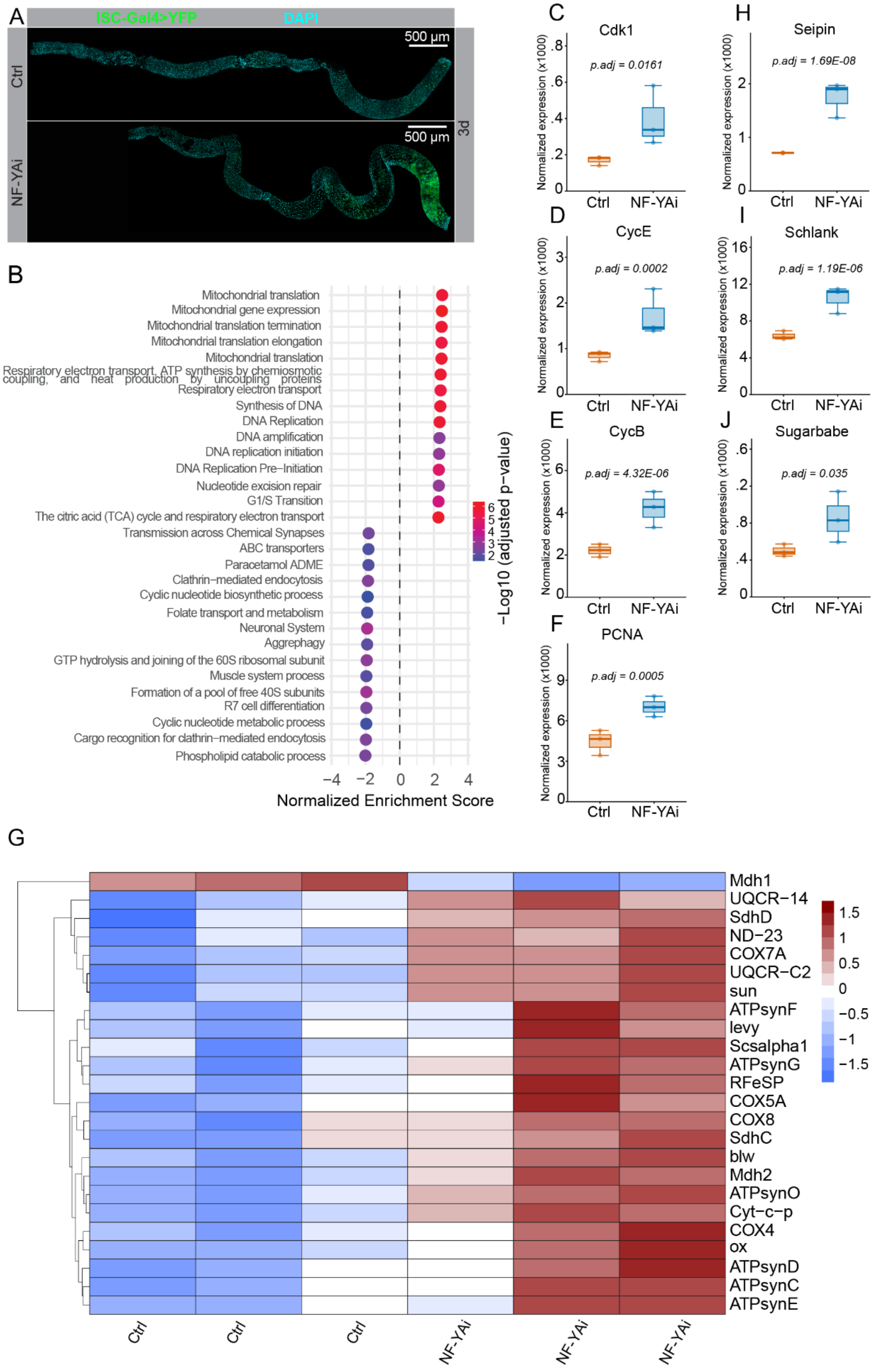


*(Legend on the next page)*

**Figure S2. Related to the main figure 3.**

(**A**) Representative images of esg-Gal4$^{ts}$, Su(H)GBE-Gal80 > NF-YA RNAi (KK) and control female midguts after 3 days at +29°C. (**B**) Gene-set enrichment analysis (GSEA) from the mRNA sequencing experiment depicted in Figure 3A. The analysis includes gene set annotations from the Gene Ontology (GO), Kyoto Encyclopedia of Genes and Genomes (KEGG), WikiPathways (WP) and Reactome databases. (**C**-**F**) Normalized mRNA expressions of *Cdk1* (C), *CycE* (D), *CycB* (E), and *PCNA* (F). (**G**) Heatmap representing differentially expressed genes of the tricarboxylic acid cycle and the electron transport chain (GO database) from the RNA sequencing experiment depicted in Figure 3A. (**H-J**) Normalized mRNA expressions of *Seipin* (H), *Schlank* (I), and *Sugarbabe* (J).

FIGURE S3

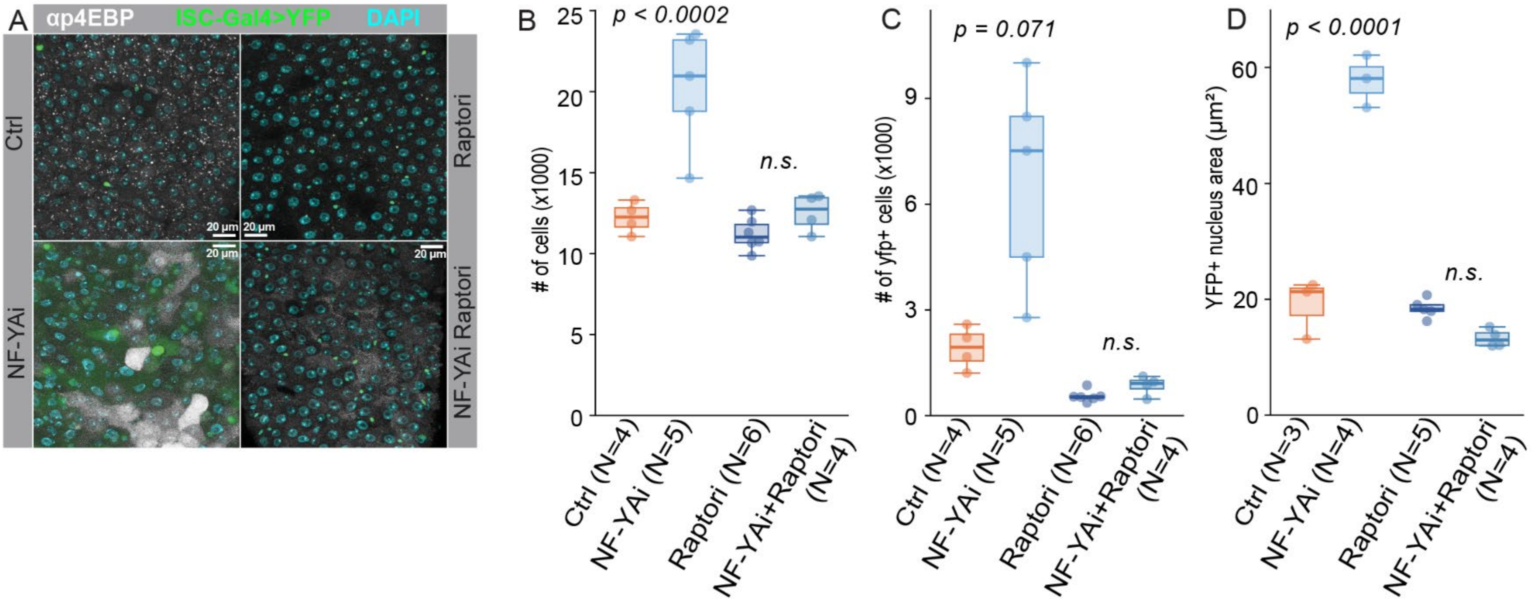


**Figure S3. Related to the main figure 4.**

(**A**) Representative images of esg-Gal4$^{ts}$, Su(H)GBE-Gal80 > NF-YA RNAi (KK), > Raptor RNAi, > Raptor RNAi + NF-YA RNAi combination, and control female midguts from the R4b region immunostained with α-p4EBP antibodies. (**B-D**) Quantification of total cell numbers (B), YFP+ cell numbers (C), and YFP+ nuclear area (D) from the experiment depicted in (A). P-value in (C) was obtained by Welch analysis of variance (ANOVA) followed by the Games-Howell test. P-values in (B) and (D) were obtained by two-way ANOVA followed by Tukey’s test.
